# Neural Dynamic Underlying Coordination Process between Habitual and Goal-Directed Behavior

**DOI:** 10.1101/2023.03.16.533062

**Authors:** Mengyang He, Wen Wen, Changzhu Qi

## Abstract

The coordination between habitual and goal-directed behaviors has significant evolutionary importance. However, the specific cognitive processes and neural mechanisms underlying this coordination process require further research. Since inducing natural habitual responses through repetitive stimuli-response training in a laboratory environment is extremely difficult in humans, well-trained sports experts with automatic perception-action features towards expertise-related stimuli serve as ideal natural samples to address this critical gap. We conducted scalp EEG recordings while sports experts performed an expertise Simon task that involved both automatic and goal-directed processes with moderate space of expertise-related stimuli congruent or incongruent with the response hand. In the congruent condition, sports experts showed larger response-locked LRP and beta band (15-25Hz) activity at the frontal-central region, indicating an enhanced automatic response tendency towards expertise-related stimuli. In the incongruent condition, a larger theta (3-8Hz) dynamic was observed in the superior frontal gyrus when sports experts needed to inhibit the automatic response tendency. The results indicated that sports experts exhibited an enhanced coordination process towards expertise-related stimuli, which was closely related to specific cognitive processes of response preparation and response inhibition in coordinating habitual and goal-directed behaviors

## Introduction

Humans typically choose their actions based on environmental information through two distinct response systems: the automatic habitual stimulus-driven response and the cognitively controlled goal-directed response. The habitual stimulus-driven response is characterized by high sensitivity to specific stimuli due to repetitive stimuli-response association training, which often leads to automatic response tendencies. The goal-directed response involves high sensitivity to current response goals, leading to the coordination of habitual behavior to achieve those goals. The coordination between these two processes is of significant evolutionary importance. However, the cognitive and neural mechanisms underlying this coordination process in the brain require further investigation.

Two main approaches (reward-induced habitation and training-induced habitation) are commonly used to induce habitual behaviors. Researchers have conducted extensive work on reward-induced habituation in both animals and humans, which has demonstrated the crucial role of reward and habituation brain networks(Gremel & Costa, 2013). However, there has been limited research on training-induced habituation in humans. The reason for this rarity is that while habitual behavior can be successfully established through stimuli-response repetition in animals, it is challenging to induce this type of behavior in a laboratory environment in humans. The complexity of human behavior and limited research time make it difficult to establish the necessary nature connection in human subjects. Cognitive and neural evidence for this process has been obtained from studies examining participants who already possess specific habitual stimuli-response connections, such as individuals with compulsive disorders and drug addiction (both of which are results of over habituation). Studies that focused on the brain activity underlying the imbalance of habitual and goal-directed control of addictive stimuli in compulsive disorders have shown significant impairment of executive brain networks(Xu et al., 2019). Other studies have examined the conflict between habituation and goals in healthy participants when habitual responses are incongruent with current goals. These studies have shown enhanced activation of executive brain networks, which are highly associated with theta oscillation(Adelhöfer & Beste, 2020). However, more direct evidence of training-induced coordination processes is still needed, and a more accurate approach to studying this coordination process in a natural environment is still lacking.

Sports experts who gained plenty of automatic perception-action features of highly habituated towards expertise-related stimuli after extensive long-term expertise training provided a perfect model to examine this issue. They showed extremely high efficiency in perception (Wang, Yang, Zhu, & Min, 2013), attention (Mengyang et al., 2018), motor preparation (Wang et al., 2013) and motor decision-making(Piras, Raffi, Perazzolo, Malagoli Lanzoni, & Squatrito, 2019; Wolf, Brlz, Scholz, Ramos-Murguialday, & Strehl, 2016) towards their expertise-related stimuli. At the meantime, sports experts also demonstrated extraordinary inhibition control ability, allow them to inhabit this habitual automatic tendency, anticipate and flexibly select appropriate response towards current environment (Nakamoto & Mori, 2012; Q. Zhao et al., 2021). This flexible coordination is achieved by both enhanced perception-action ability of and stronger habitual process inhibition control ability of adaptive behavior. Hence, sports experts become a perfect sample to explore the coordination between habitual behavior and inhibition control of this habitual behavior in nature environment.

Previous studies have demonstrated some EEG evidence of single process for this coordination process of sports experts. For the habitual process, sports experts showed significant automatic respond tendency towards expertise-related stimuli, this enhanced stimuli-response effect was consistent with a larger CNV and Alpha amplitude. For the adaptive process, the enhanced ability of inhibition control for sports experts is linked to greater frontal P300s and larger amplitude N2s compared to controls (Muraskin, Sherwin, & Sajda, 2015; Nakamoto & Mori, 2012). However, how sports experts coordinate these two processes towards expertise-related stimuli and the neurobiological mechanism underlying this process needs to be verified with evidence from more complex tasks, which engage the detection of competing responses and the selective inhibition of inappropriate responses.

Simon task has been widely used to examine the goal-directed process which needs to inhibit cognitive interference that occurs when processing a specific stimulus whose space is incongruent with the response. The logic of Simon task is that the congruency of the stimuli and response hand could indue an automatic habitual response tendency (Simon, 1969), thus there would be a larger conflict in incongruent trials compared to congruent trials. The present study explored the training-induced coordination by developing an expertise-related Simon task to examine how sports experts coordinate between the automatic habitual tendency and adaptive goal-directed process towards their expertise-related stimuli. In the present study, we combined Simon task with expertise-related stimuli, thus, the paradigm includes two different conditions: stimuli-response congruence (referred as expertise-related stimuli congruent with the response hand) and stimuli-response incongruence(referred as expertise-related stimuli incongruent with the response hand). Based on the automatic response activation effect, the expertise-related stimuli would evoke a stronger automatic response tendency in stimuli-response congruence condition. Correspondingly, a stronger inhibition control effect would be evoked in the condition of stimuli-response incongruence.

To this end, we recorded scalp EEG of sports experts when they were performing the expertise-Simon task to examine the neural mechanism underlying this coordination process. In addition to the behavioral features including reaction time and accuracy rate, we hypothesis that sports experts would induce stronger LRP reflecting response process and an typical component reflecting inhabitation control(Dambacher et al., 2014; Mückschel, Dippel, & Beste, 2017).The further purpose in the present study is to trace the brain activation involved in the inhabitation control, as the expertise-relevant stimuli induce more response inhabitation than expertise-irrelevant stimuli.

## Methods

### Participants

A prior power analysis suggested that a sample size of twenty-four would yield a medium effect size (d = .25) and a high statistical power (80%) given a two-by-two repeated design (alpha =.05). We tested twenty-six table tennis players of the provincial team in case of drop out and exclusion. The inclusion criteria were : (1) Skill level: qualified as the National Player at or above First Grade; (2) Training experience: more than 10-year experience in professional training; (3) Training frequency: playing table tennis more than 5 times per week and above 8h each time. The final sample had twenty-four participants (mean age =18.1, 14 males, all right-handed) since two participants have less than 70% valid trials after preprocessing. There was no significant difference in age (male: 18.2±0.30, female. 18.1±0.31, t(1,22)=0.26, p=0.799) nor training years (male: 10.1±0.29, female, 10.4±0.62, t(1,22) =-0.53, P = 0.601) between male and female players. All participants had normal or corrected-to-normal vision, and they reported no history of neurological or psychiatric disorders. Written consent forms were obtained from all participants and they received remuneration as a compensation. This study was approved by the Institutional Research Ethics Committee of Wuhan Sports University (2022008).

### Stimuli and Design

Participants sat in front of a LED monitor at a distance of 70cm and were presented with different images (6.23°×6.23°) on a gray background. Stimuli were presented using the Psychtoolbox-3(Brainard, 1997) in MATLAB 2018b (Mathworks, MA, USA). There were two types of stimuli: the expertise-relevant stimuli (four images) contained a table tennis racket and a table tennis ball whereas the neutral stimuli (four images) contained a baseball glove and a baseball. All images were standardized using the SHINE software to calibrate the grayscale (hue, lightness and chroma) and size(Willenbockel et al., 2010) (Li, Zhang, Liu, & Luo, 2022). Each trial started with a black fixation cross (0.5°×0.5°) displayed at the center of the screen. After a jittered time interval (500-700ms), one image would be presented at either the left or the right side (10° to the center) with equal probabilities and lasted for 300ms. Participants were instructed to respond to the type of stimuli using the corresponding hand within 2 seconds. For instance, the participant in the Figure 1 was instructed to indicate the expertise/neutral stimuli with right/left hand (by pressing “m/z” on the keyboard). Based on the relationship between stimuli location and response hand, we categorized the trials into congruent and incongruent conditions. Figure 1A concluded two different conditions, the congruent condition where the expertise image is presented on the right side that is compatible with the response hand; the incongruent condition where the neutral image is presented on the right side that is incompatible with the response hand. The stimuli-response mapping was counter-balanced between participants. Therefore, there were four conditions with the combination of image type and location-response congruency effect: expertise congruent condition (EC), expertise incongruent condition (EIC), neutral congruent condition (NC) and neutral congruent condition (NIC). Each condition had 120 trials, so there were 480 trials in total, trials from different conditions were mixed in randomized order within each block.

**Figure 1.**
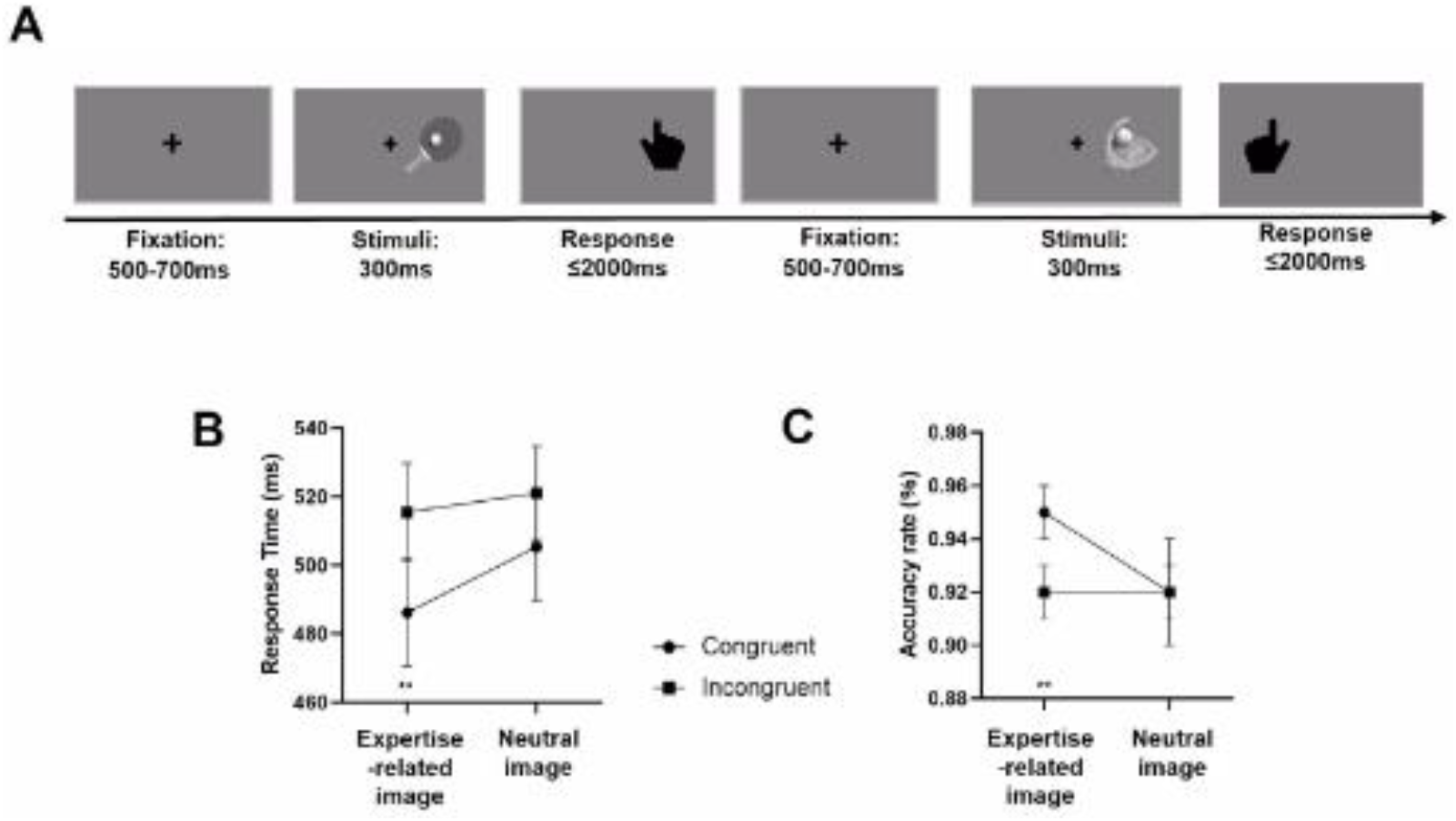
(A)Expertise-related Simon task. Participants were instructed to indicate expertise-related or neutral stimuli using the right or left index finger.. From left to right represent different trials of Examples illustrate the expertise congruent trial (Con E: The location of the expertise-realted stimuli was congruent with the correct response hand) and the Neutral Incongruent trial (: the location of the netural stimuli was incongruent with the correct response hand). (B)&(C) Behavioral performance: (B)mean reaction times (RT) and (C)accuracy of each condition. errorbar represent CI or standard error? p<.05, *, p<.01, **p<.001, ***error bar represents

**Figure 2.**
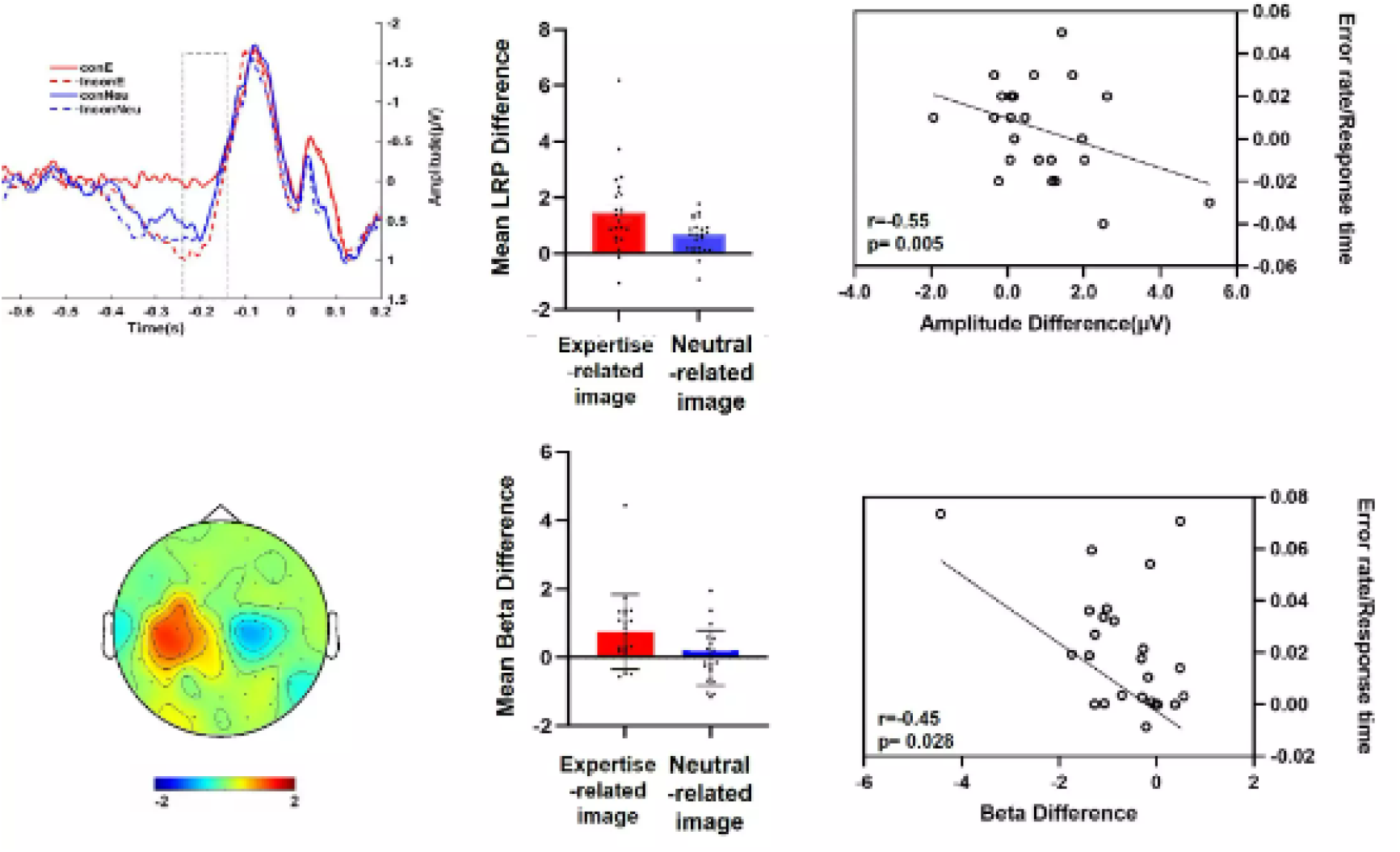
A represents the results of image-decoding. The shaded area represents the standard deviation of 5000 times bootstrapped mean. The solid/dashed vertical line indicated the image onset/offset. Bold bar below the x-axis indicated significant time-clusters. B represents the response related LRP difference. Each dot correponds to individual’s data. C Scatter plots (with best-fitting regression lines) illustrates the correlation between amplitude difference as a function of the RT (right panel).

### EEG Recording and Preprocessing

Electroencephalogram (EEG) data was recorded from Ag/AgCl electrodes placed according to the international 10-20 system with BrainAmp amplifier (Brain Product). Ground electrode was FPz. All electrodes were online referenced to FCz. Impedance between electrodes and scalp was keep below 5 KΩ. The data were collected at a sampling rate of 1000 Hz and band-pass filtered to 0.1–100 Hz. Offline data analysis was performed using EEGLAB (Delorme and Makeig, 2004). Raw data were first referenced to the average of all electrodes and then band pass filtered to 0.1–40 Hz with a slope of 24db/Oct. Data was epoched between −1000 and 2000ms relative to the onset of the stimuli and the average baseline was removed using data from - 200ms to 0ms. Independent Component Analysis was used to detect and correct eye blinks, eye movements, heart and line noise. Finally, epochs containing voltage deviations exceeding ±50 μV or contaminated by muscular artifacts were excluded by visual inspection. There were 96.9% clean trials after preprocessing.

### Behavioral Data

Trials with RTs beyond three standard deviations of mean were excluded from the analysis. Mean RT of remaining correct trials in each condition was then computed. Accuracy was the proportion of the number of correct trials against the total number of trials in each condition. A two-by-two repeated ANOVA was performed on RT and accuracy with stimuli type (expertise vs. neutral) and location-response congruency (congruent vs. incongruent) as the within-subject factors.

### Event-Related Potential Analysis

For stimulus-evoked lateralized readiness potentials ( LRP), data was re-epoched to −200 to 800 ms relative to stimulus onset and baseline corrected to the pre-stimulus interval. For response-evoked LRP, data was re-epoched to −800 to 200 ms relative to response onset and baseline corrected to the interval of −1000 to 800 ms relative to response onset. LRP was calculated as the difference wave between contralateral and ipsilateral channels. Central electrode C3 and C4 were used for assessment of LRP as suggested by previous studies(Kyung Hun Jung, Martin, & Ruthruff, 2021).For instance, if the stimuli was required to respond by left hand, LRP was calculated using C4 minus C3. A two-by-two repeated ANOVA(stimuli type: expertise vs. non-expertise; spatial congruency: expert vs. novice) was conducted on the peak amplitude of the LRP.

The peak amplitude of stimulus- and response-locked LRP was identified at the group level in each of the 4 conditions. For stimulus-locked LRP, the peak detection started from 200 ms post-stimulus onset, the peak detection was conducted through the time window of The mean amplitude was then calculated by averaging 50 points (100 ms) centered at the peak point. For response-locked LRP, the peak detection started from response onset and went backwards, the peak detection was conducted through the time window of −250—-150ms. The mean amplitude was calculated by averaging 50 points centered at the group peak point.

### Time Frequency Analysis

Time-frequency analysis was conducted using Matlab based Fieldtrip toolbox (Oostenveld et al., 2011). EEG data were performed using FFT functions of single-trial data prior to averaging. Spectral amplitudes for each frequency were estimated for the range from 1 to 40 Hz, in 2-Hz steps, during the time window of −500 to 1500 ms relative to stimulus onset in steps of 20 ms. The theta-band (3-8 Hz) and beta (13-20Hz) band were extracted for further analysis. This time-frequency analysis was performed for each condition and for each subject separately, the grand mean of time-frequency power was then averaged across subjects. The theta-band power profiles were normalized by subtracting the averaged - 200-0 theta-band time courses from each theta -band and in each subject respectively. Given the prominent theta-band activations in the inhibition control, we selected frontal-central 9 channels (F1, F2, Fz, FCz, FC1, FC2, Cz, C1, C2) for further analyze. Given the prominent beta-band activations of motion action, central electrode C3 and C4 were used for assessment of the mean amplitude of beta as suggested by previous studies() for further analyze. The beta-band power profiles were then calculated by subtracting contralateral channel form ipsilateral channels. For instance, if the stimuli was required to respond by left hand, beta amplitude was calculated using C4 minus C3.

### Source localization

Standard Low-Resolution Electromagnetic Tomography (sLORETA) was used to localize cortical electrical activity of inhibition control. LORETA estimates the distribution of electrical neural activity based on the measurements of a dense grid of scalp electrodes. sLORETA (Pascual-Marqui, 2002), estimates the current source density distribution for epochs of brain electrical activity on a dense grid of 6239 voxels at 5 mm spatial resolution. We constructed a three-shell spherical model of the brain for each subject. Lead fields were calculated based on electrodes and head model. The Simon effects on sLORETA were compared between groups with t-statistical non-parametric mapping, using the implemented statistical nonparametric mapping (SnPM) tool. The significance level applied to the data was set at pb0.05 (significant effect) and pb0.10 (statistical trend). To improve inter-subject comparability, MNI space data was smoothed with a Gaussian kernel with smoothing parameter value of 0.5. Statistical comparisons between source locations were undertaken using an implementation of Statistical Non-Parametric Modelling.

### Decoding

Mahalanobis decoding was performed based on the whole-brain electrodes’ broad-band EEG data. Data re-epoched into −0.2s~1.6s relative to the stimulus onset and baseline-corrected to the mean amplitude of the pre-stimulus period. We gaussian-smoothed (window size = 16 ms) the data along the time dimension to reduce temporal noises. To avoid the confounding of the stimulus location, we performed decoding separately for trials where stimulus appeared on left/right and used the averaged results for statistical analysis. At each time point, data was partitioned into the seven training-fold and one test-fold. To create an unbiased classifier, the number of trials for each type in the training set was equalized by subsampling. The covariance matrix was estimated based on the trial-averaged training set. Pairwise Mahalanobis distances between test-trials and the averaged training data were computed using the covariance matrix. A smaller distance toward the activation profile of one stimuli type indicated a larger pattern similarity. Therefore, the decoding was marked as a success when the distance between this given trial and its corresponding activation profile in the training data was smaller than the other stimuli type. Cross-validation was carried out in a leave-one-fold-out manner. This routine was iterated 1000 times and trials were randomly assigned to the training and test set in each iteration. Further statistical analysis was performed on the averaged decoding accuracy. After gaussian-smoothing (window size = 16 ms), the accuracy time series were compared to the chance level (1/2). Cluster-based permutation was performed on the decoding accuracy time-series at the group level (alpha = 0.05, cluster-based nonparametric alpha = 0.05, cluster 0statistic = sum, two-tail, permutation times = 10000).

## Results

### Behavior : Experts showed larger congruency effect on expertise-related stimuli

There was a significant main effect of stimuli (expertise vs. neutral), F(1, 23) = 7.55, p =0.011, η2p = 0.247, revealing that RTs for the expertise-relevant stimuli (500ms) were faster than that for the neutral stimuli (513ms). participants were faster in congruent trials (495ms) than in incongruent trials (518ms, F(1, 23) = 30.83, P < 0.001, η2p= 0.573). Moreover, the significant interaction between expertise and congruency (F(1, 23) = 8.08, P =0.009, η2p = 0.260) driven by larger congruency effect of expertise stimuli (29ms, F(1, 23) = 11.1, p = 0.003) than neutral stimuli(16ms, F(1, 23) =0.07, p = 0.789) suggested that sports experts manifested stronger automized response towards expertise-relevant stimuli.

Analyses on accuracy revealed a significant main effect of congruency (F(1, 23) = 4.52, P = 0.044, η2p = 0.164). There was no significant main effect of stimuli type(F(1, 23) = 1.35, P = 0.258, η2p = 0.055), nor the interaction effect (F(1, 23) = 2.27, P = 0.145, η2p = 0.164). However, sports experts showed larger accuracy rate of expertise stimuli (0.96, F(1, 23) = 5.59, p = 0.027) than neutral stimuli(0.92, F(1,23) =0.2, p = 0.661).

### Perception-action feature of habitual response process towards expertise-related stimuli for sports experts

#### 1) Sports experts showed enhanced sensitivity during the perception of expertise-related stimuli

We first performed Mahalanobios-distance decoding to test whether the two types of stimuli evoked distinctive neural activation patterns in early perception stage. As shown in Fig 3, experts could discriminate expertise-relevant and neutral images right over stimulus-onset (54~868ms, clustered p <.001) indicating enhanced perceptual sensitivity for expertise-relevant stimuli. This supported our claim that automatic processing emerged at the perceptual level.

**Figure 3.**
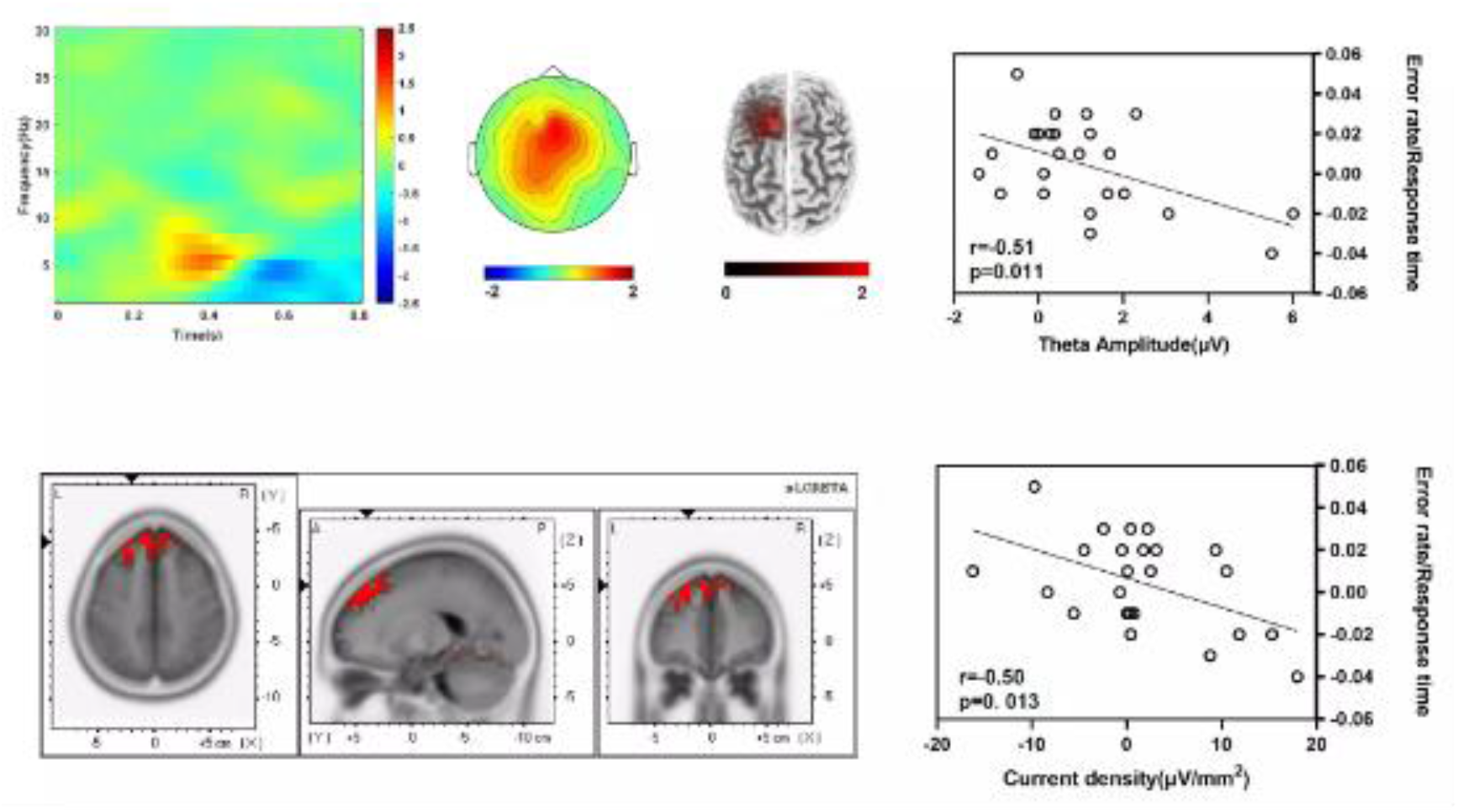
Frontal theta. (A) (B) (C)Topographical map of theta activity. The averaged theta activity during the 350~500ms interval. A topographical distribution of frontal theta activity (350~500ms). B averaged theta power of each experiment condition. black dots correspond to each individual. error bars represent SE. C. source localization of theta activity for sLoreta.

#### 2) Sports experts showed enhanced ability of response preparation towards expertise-related stimuli

The peak amplitude of stimuli-locked LRP showed no significant difference between congruency and incongruent trials.

The peak amplitude of response-locked LRP showed a significant difference between congruency and incongruent trials, F(1, 23) = 32.93, p <.001, η2p = 0.589, with more negative LRP in the congruent conditions than in incongruent conditions. The main effect of group was not significant, (1, 23) = 2.33, p =.14, η2p = 0.092. The interaction between expertise and congruency was significant, F(1,23) = 9.07, p < 0.001, η2p = 0.283, which was driven by a larger amplitude difference between EIC trials and EC trials(EC vs. EIC, 1.47, p=0.07) than the difference between NIC trials and NC trials (NC vs. NIC,0.6, p=0.281). Figure 3B shows the averaged waveforms of the response-locked LRP. The amplitude difference modulated by expertise showed a correlation with RT difference modulated by expertise, r =-0.55, p =0.005.

Next, we examined response-locked beta-band activity over central regions, which is an index for motor action. The main effect of stimuli was significant, the amplitude of central beta was stronger in congruent than incongruent trials, F(1, 23) = 12.66, p =.002, η2p = 0.355. The main effect of group was insignificant, F(1, 23) = 0.83, p =.372, η2p = 0.035. The interaction between expertise and congruency was significant, F(1,23) = 4.56, p =.044, η2p = 0.166, which was driven by a larger amplitude difference between EC trials and EIC trials(EC vs. EIC, 0.71, p=0.003) than the difference between NC trials and NIC trials(NC vs. NIC,0.05, p=0.738).The amplitude difference modulated by reward showed a correlation with RT difference modulated by expertise, r =-0.45, p =0.028.

### Experts showed stronger response inhibition for expertise-relevant stimuli

The topographical distribution of theta band oscillations is shown in Figure3. Midfrontal cluster were selected with the frequency ranging from 3 to 8 Hz and the time interval ranging from 350 to 500 ms post onset. The 2×2 ANOVA on theta oscillation showed a main effect of stimuli type and congruence, which revealed a significant main effect of congruence, F(1,22)=4.42, p = 0.05,η2p=0.17,with larger brain activity to the expertise-relevant stimuli than non-expertise-relevant stimuli. The interaction between stimuli type and spatial congruency also significant, F(1,22) = 8.56, p = 0.01, η2p = 0.28, which was driven by a larger amplitude difference between EIC trials and EC trials (EC vs. EIC, 1.26, p=0.07) than the difference between NIC trials and NC trials (EC vs. EIC, 0.12, p=0.07). The amplitude difference modulated by reward showed a correlation with RT difference modulated by expertise, r =, P =.

The superior frontal gyrus was chosen as ROI since it is a classic area for response inhibition. The 2×2 ANOVA on current density showed a main effect of congruence, F(1,22)=9.73, p = 0.005,η2p=0.297,with larger brain activity to the expertise-relevant stimuli than non-expertise-relevant stimuli. The interaction between stimuli type and spatial congruency also significant, F(1,22) = 7.40, p = 0.12, η2p = 0.24, which was driven by a larger amplitude difference between EIC trials and EC trials (EC vs. EIC, 10.83, p=0.07) than the difference between NIC trials and NC trials (EC vs. EIC, 2.05, p=0.07). The difference of current density modulated by expertise showed a correlation with RT difference modulated by expertise, r = 0.569, p = 0.004.

## Discussion

The present study was designed to examine the cognitive neural mechanism underlying the coordination of training-induced habitual and goal-directed process using an expertise-Simon task.

### Enhanced perception-action process for expertise-related stimuli revealed by Simon task

Here, we found a lager Simon effect for expertise-related stimuli than for neutral stimuli. the difference of Simon effect between expertise-related stimuli and neutral stimuli reflects the enhanced cognitive process underlying this coordination process. The expertise-relevant stimuli would evoke stronger automatic habitual response tendency towards expertise-related stimuli compared to neutral stimuli in congruent condition and enhanced inhabitation control of in incongruent condition. The results were consistent with the previous research related to sport experts’ perception-action coupling which suggested that sport experts had a faster response time among baseball-specific stimuli (Nakamoto and Mori, 2008). Taken together, sports experts showed a significant larger coordination effect of expertise-related stimuli. The whole coordination process consists of the enhanced automatic perception-action features for habitual process and inhibition control features for goal-directed process.

### Role of response preparation for expertise-related stimuli of sports experts

Our study provided further evidence to support the enhanced habitual process, as demonstrated by typical EEG elements. Specifically, our results showed that expertise-relevant stimuli induced a stronger response-locked LRP, which originates from the motor system and represents response preparation. (Chen, van den Berg, & Kwak, 2022; Takacs, Bluschke, Kleimaker, Münchau, & Beste, 2021), K. H. Jung, Martin, & Ruthruff, 2020; Osman, Moore, & Ulrich, 2003), This finding suggests enhanced motor activation by expertise-relevant stimuli. Additionally, consistent with this result, expertise-relevant stimuli also induced a stronger response-locked beta oscillation, which is typically associated with motor preparation. Prior research has shown that beta oscillatory activity (13-20 Hz) is present during motor preparation, (Little, Bonaiuto, Barnes, & Bestmann, 2019). Particularly during the ERD period., The timing of beta bursts has been linked to the degree of motor preparation. (Little, Bonaiuto, Barnes, & Bestmann, 2019). These results are consistent with previous studies that have examined the specific cognitive abilities involved in this coordination process by analyzing the time-dependent competition between goal-directed and habitual trails. These studies have revealed that response preparation, as a crucial feature of perception-action, plays an important role in this process.

### Role of response inhabitation for expertise-related stimuli of sports experts

In the human brain, the role of frontal theta band oscillations dynamic by the superior frontal gyrus during response inhibition is very clear(Adelhöfer, Mückschel, Teufert, Ziemssen, & Beste, 2019). Previous research explored the neural mechanism underlying goal-directed behavior and the particularly demand to inhibit automatically executed responses(Adelhöfer et al., 2019; Pscherer, Mückschel, Summerer, Bluschke, & Beste, 2019).The eeg and eye data showed that inhibitory control processes reflected by theta oscillations are strongly modulated by the neural processes in the <u>SMA</u> and <u>SFG</u> (Dippel, Mückschel, Ziemssen, & Beste, 2017). Along these lines, our results further revealed that response inhibition by superior frontal theta could be modulated by sports expertise. These results are consistent with the study that examined how the sports experts process congruency monitoring while observing expertise-relevant actions. For the halfpipe trick, experts exhibited better task performance and greater midline theta oscillations before possible incongruency compared with controls. Source reconstruction for the theta oscillation revealed a stronger activation in the middle and superior frontal gyrus for experts in response to incongruency compared with controls (Qiwei Zhao et al., 2021). Standard Low-resolution brain electromagnetic tomography is an instantaneous, distributed, discrete, and linear EEG/MEG inverse solutions, yields images of standardized current density with zero localization error(Boughariou et al., 2015; Pascual-Marqui, 2002). The technique has been successfully used to reveal temporal effect of scalp EEG power spectrum in cognitive tasks(Imperatori et al., 2014; Imperatori et al., 2013).

In conclusion, sports experts showed enhanced coordination process towards expertise-related stimuli, this process was specifically related to motor preparation and motor control.

## Notes

### Competing Interest Statement

The authors have declared no competing interest.

